# Comparative analysis of the EF-1α Intergenic Region in *Babesia divergens* isolates: Insights into TA Repeat Variation and Potential Regulatory Implications

**DOI:** 10.1101/2025.08.08.669274

**Authors:** Sezayi Ozubek, Alejandro Sanchez-Flores, Estrella Montero, Heba Alzan, Carlos E. Suarez, Ricardo Grande, Aitor Gil-Iglesias, Munir Aktas, Luis Miguel Gonzalez

## Abstract

*Babesia divergens*, a zoonotic tick-borne pathogen, causes bovine and human babesiosis in Europe. The Elongation Factor 1 alpha (EF-1α) protein is important in many cellular processes and has emerged as a possible target for subunit vaccine development against parasitic infections, and its intergenic region (IG) is an important tool for genetic manipulation of *Babesia* parasites. While the EF-1α locus of *B. divergens* has been described, structural variation between isolates was poorly defined. In order to fill this gap, we performed a comparative analysis of the EF-1α-IG region in *B. divergens* human (Rouen 87, and Spanish sample), and bovine (Turkiye) host isolates. Our findings revealed both conserved and variable elements, particularly in TA nucleotide repeat numbers and IG sequence length. The Spanish isolate exhibited the highest TA repeat expansion, whereas the Rouen 87 strain had the shortest IG region. Given the known role of repeat-rich promoter elements in gene regulation, these differences may influence EF-1α transcription. Additionally, these findings provide insights into the evolutionary divergence of *B. divergens* and its host adaptation mechanisms. This study establishes a foundation for future gene editing and transfection strategies, where selecting intergenic sequences with varying TA repeats could optimize transfection efficiency and explain phenotypic differences between isolates from different hosts or regions.

## Introduction

Elongation factor 1 alpha (EF-1α) is among the most abundant proteins ever reported to be expressed by eukaryotic cells and is highly conserved [1,2,3]. Traditionally, EF-1α is chiefly acknowledged for its contribution to the translation machinery, facilitating the binding of aminoacyl-tRNA to the ribosomal A site in a GTP-dependent manner [4]. Nevertheless, recent studies have revealed how it functions in various cellular processes such as microtubule severing, ubiquitin dependent proteolysis of N-terminal blocked proteins and cytoskeletal rearrangements [3,5]. In addition, EF-1α plays a crucial role in fundamental cellular events including of programmed apoptosis in which is up-regulated [3,6,7]. Further studies have also highlighted the important role of EF-1α in other apicomplexan parasites, including *Cryptosporidium parvum* and *Toxoplasma gondii*, for the entry process of these parasites into host cells. This suggest that EF-1α could be a promising target for vaccine development against cryptosporidiosis and toxoplasmosis [8,9]. Furthermore, results from studies on vaccination against avian coccidiosis indicate that EF-1α is a promising candidate antigen for eliciting cross-protective immunity against this disease in poultry, highlighting its potential for veterinary use [10].

*Babesia* are *T. gondii* and *Plasmodium*-related apicomplexan tick-borne parasites responsible for acute and persistent infections in vertebrate hosts, including cattle and humans. Advancements in gene editing and transfection systems are essential for unraveling the molecular intricacies of *Babesia* parasites and enhancing vaccine development strategies [11,12,13]. Among the crucial components, facilitating these advancements is the EF-1α, which plays a pivotal role in various cellular processes. In both *Plasmodium* and *Babesia* parasites, the *EF-1α* gene exhibits a distinctive organization, featuring two identical head-to-head genes, each separated by intergenic regions [11,14]. The EF*-*1α intergenic region (EF-1α-IG) hosts robust promoters capable of driving the expression of both *EF-1α* genes efficiently. This unique characteristic of the EF-1α-IG region also enables it to effectively promote the expression of foreign genes in transiently and stably transfected *Babesia bovis* and *Babesia bigemina* parasites [11,12,15]. The utilization of these strong promoters within the EF-1α-IG region has dramatically accelerated the development of transfection and gene editing systems in both *Plasmodium* and *Babesia* parasites [11,16]. Researchers have successfully exploited these promoters to introduce foreign genes into parasite genomes, facilitating precise genetic manipulations, the development of novel vaccines, and sophisticated studies on parasite biology. This breakthrough has significantly improved gene function analysis, advanced our understanding of the molecular mechanisms underlying parasite pathogenesis and host-parasite interactions [13,17,18], and became an important tool for developing novel vectored vaccines [19,20,21,22].

*Babesia divergens*, one of the three major species of *Babesia* causing babesiosis in cattle, is particularly significant as a zoonotic pathogen playing a role in human babesiosis in Europe, in addition to its impact on cattle [23,24]. Compared to *B. bovis* and *B. bigemina*, limited information is available about the infectivity and pathogenicity of *B. divergens*, leaving many questions unanswered regarding important processes. Understanding the gaps of knowledge on gene function and other mechanisms involved in the biology of *B. divergens* will likely require the identification of novel promoters that can aid in the development of gene editing and transfection systems. Thus far, only one study has drawn attention in this regard, describing a stable transfection system for *B. divergens* based on the promoter of the *elongation factor Tu GTP binding domain family protein* (*ef-tgtp*) gene [25]. Unlike other *Babesia* species found in cattle, *B. divergens* has a broader host spectrum, including humans. We hypothesize that the EF-1α locus of *B. divergens,* a parasite able to infect bovine and human hosts, exhibits structural and regulatory variations among different isolates, which may be linked to its host adaptability, evolutionary divergence, and geographic distribution. Investigating these differences will enhance our understanding of *B. divergens* biology and provide insights into potential mechanisms of gene regulation and parasite-host interactions.

Here, based on existing genome data and NGS and Sanger sequencing, we focused on conducting a comparative analysis of the EF-1α locus from *B. divergens* isolates obtained from humans and cattle in different geographical regions, using the *B. divergens* Rouen 87 (R87) genome as a reference [26,27]. Our approach focused on conducting a comparative analysis of the EF-1α locus, which serves as a pivotal component in transfection investigations, among *B. divergens* isolates sourced from infected bovines in Turkiye and from patients with severe babesiosis in Spain [28]. The data generated in this study implies the possible influence of evolutionary selective forces shaping key sequence and structural differences in the EF-1α locus between these two isolates.

## Results

### EF-1α locus organization and synteny in *Babesia divergens*

In apicomplexan parasites, the *EF-1α* gene locus predominantly resides between the *Ribonucleoside diphosphate reductase* and *Glutamyl tRNA* genes as demonstrated by conserved synteny between *Theileria* and *Babesia* species. Similar to *B. bovis*, *B. bigemina*, *Babesia* sp. Xinjiang and *Theileria equi*, the *B. divergens* R87 and Spanish and Turkiye isolates also possess two identical *EF-1α* genes located within the EF-1α locus (Figure 1). The biological significance of remarkable conserved synteny among all these apicomplexan parasites remains uninvestigated.

**Figure 1:**
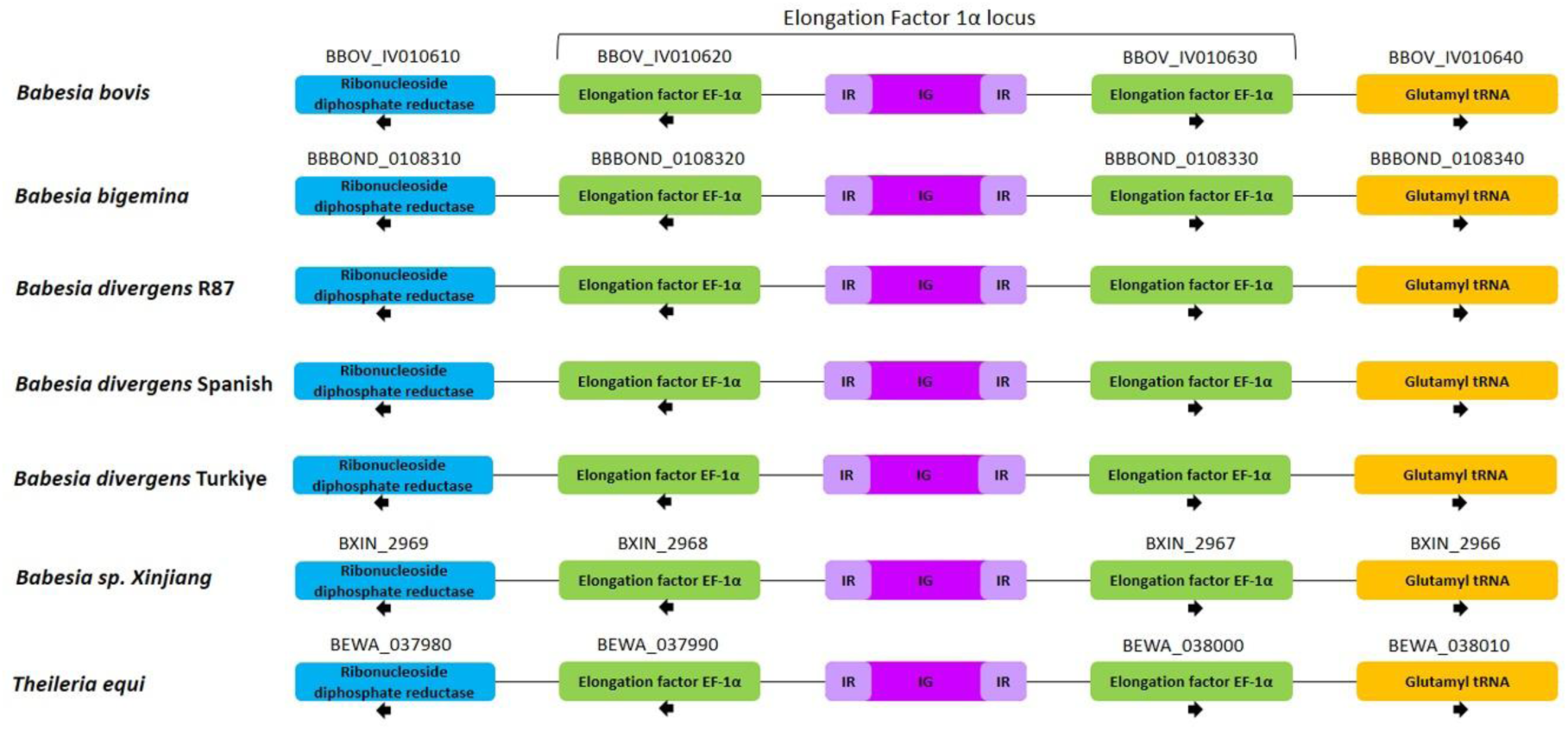
Schematic representation of gene localization and synteny maps of the *EF-1α* gene locus.

### EF-1α coding sequence conservation across *Babesia divergens* isolates

Comparative analysis of the EF-1α locus in *B. divergens* isolates (R87, Spanish, and Turkiye isolates) revealed that the *EF-1α* gene sequences (1347 bp) were 100% identical across all isolates. Each of the two EF-1α ORFs from three isolates of *B. divergens* encodes identical proteins of 448 amino acids, with an estimated molecular mass of 49.39 kDa, which is the same size as those encoded by *B. bovis* (XP_001610983.1), *B. bigemina* (XP_012766720.1), *Babesia caballi* (GIX62688.1), *Babesia gibsoni* (KAK1443338.1), *Babesia duncani* (KAK2195768.1), *Babesia ovis* (GFE54231.1), *Babesia* sp. Xinjiang (XP_028869924.1), *Theileria parva* (XP_766247.1), *Theileria annulata* (XP_954051.1), and *Theileria orientalis* (UKK00547.1). The EF-1α *B. divergens* R87, Spanish, and Turkiye ORFs show identical sequences (100% identity) and share significant similarity with homologous sequences found in *B. gibsoni* (97.99% identity), *B. caballi* (97.54% identity), *B. bigemina* (96.65% identity), *Babesia* sp. Xinjiang (96.43% identity), *B. bovis* (94.87% identity), *B. ovis* (94.20% identity), *B. duncani* (93.97% identity), *T. orientalis* (92.41% identity), *T. parva* (89.51% identity), *T. annulata* (89.06% identity), and *B. microti* (86.58% identity) (Supplementary Figure 1). Notably, key regions involved in guanosine 50-triphosphate (GTP) binding are conserved across these organisms, as depicted in Supplementary Figure 2. A phylogenetic analysis using the EF-1α protein sequence of R87, Spanish, and Turkiye isolates, along with other related piroplasma species, showed that both isolates clustered within the same clade as *B. gibsoni* (Figure 2a). Similarly, in the phylogenetic analysis of *B. divergens* R87, Spanish (MG944238.1), and Turkiye (PP663285.1) *18S rRNA* gene sequences, it was observed that they were in the same clade with *B. divergens* isolates obtained from different hosts from GenBank (Figure 2b).

**Figure 2:**
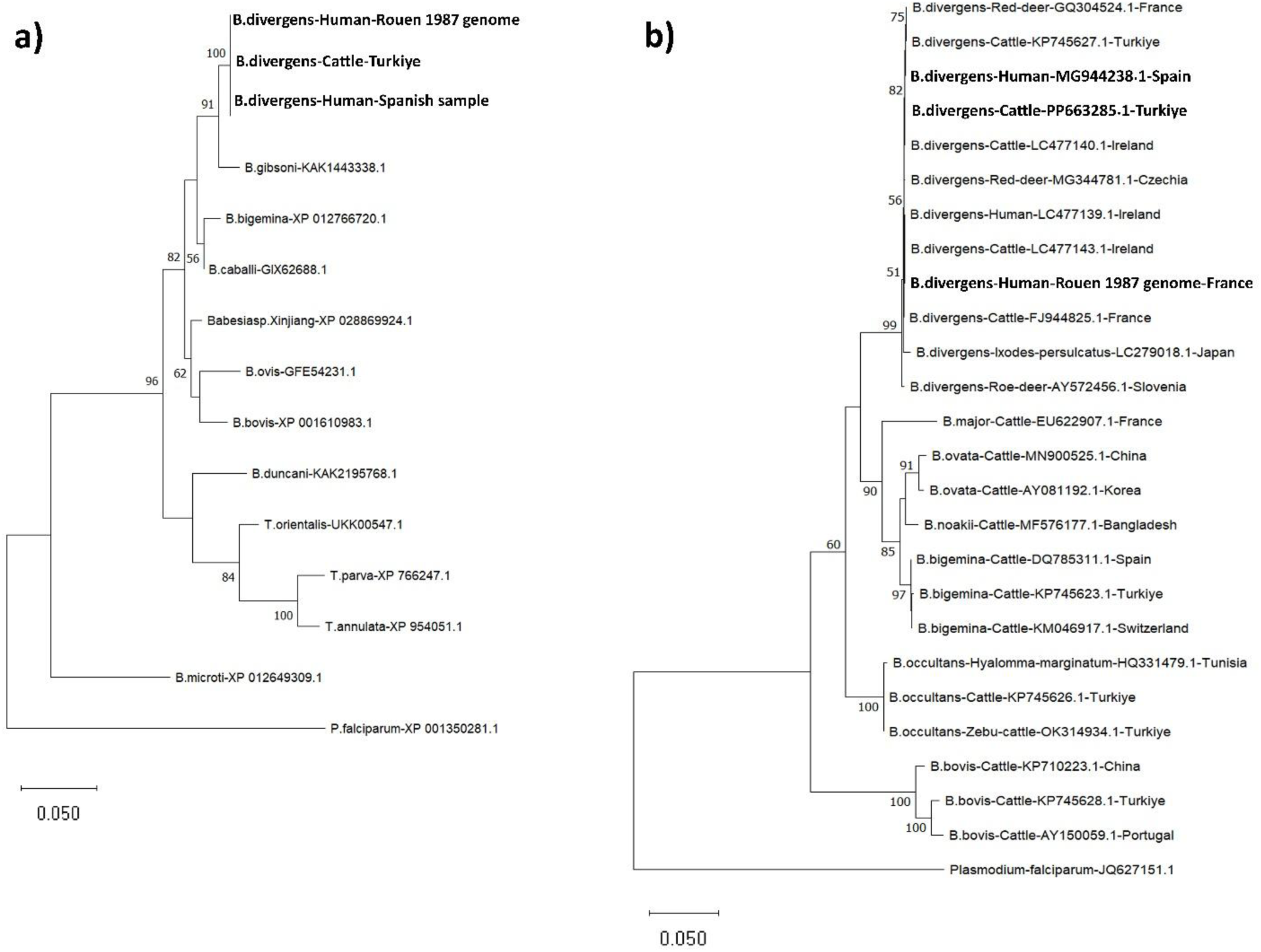
a) Phylogenetic analyses of piroplasm EF-1α protein sequences by maximum likelihood method based on the Le_Gascuel_2008 model [29]. Sequences were obtained from the GenBank database. The EF-1α protein sequence of *Plasmodium falciparum* was utilized as an outgroup. b) Phylogenetic analyses of piroplasm *18S rRNA* gene sequences by maximum likelihood method based on the Tamura-Nei model [30]. The *18S rRNA* gene sequence of *P. falciparum* was utilized as an outgroup. *Babesia divergens* R87, Spanish, and Turkiye isolates of EF-1α are highlighted in bold.

### Features, sequence alignment and nucleotide variability in IG region

The EF-1α-IG regions of the three distinct *B. divergens* isolates analyzed hereby are not related in sequence to the IG regions of other related *Babesia* parasites. However, the *B. divergens* EF-1α-IG regions do have structural similarities with other *Babesia* parasites, including the presence of terminal inverted repeat (IR) regions of 237 bp, which were conserved among all isolates. In addition, similar to what was found on *B. bovis* and *B. bigemina* [11,12], the IR regions harbor a 153 bp intron, maintaining sequence integrity across isolates (Figure 3).

**Figure 3:**
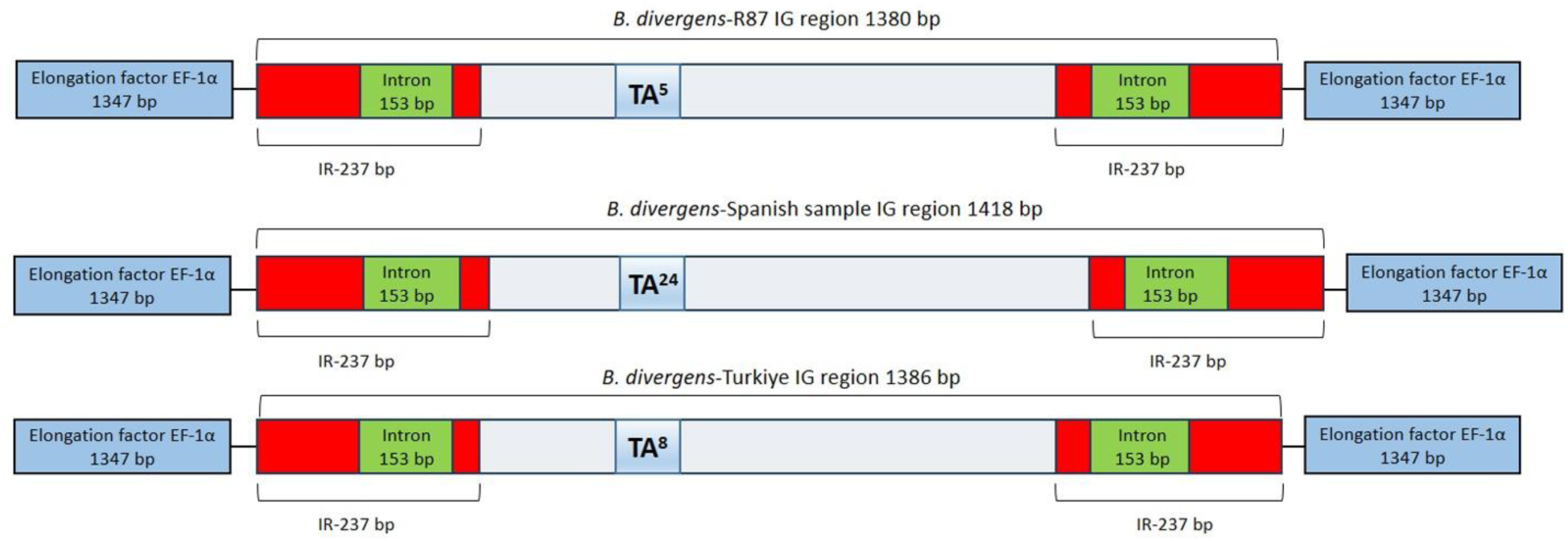
Schematic representation of the EF-1α-IG region between *B. divergens* isolates. The terminal inverted repeat (IR) regions (red), intron (green, 153 bp), and TA repeats (blue) are highlighted. The IG region length varies due to differences in TA repeat numbers, with 5 TA repeats in R87 (1380 bp), 24 in the Spanish isolate (1418 bp), and 8 in the Turkiye isolate (1386 bp).

However, the comparative analysis of the EF-1α locus sequences in the three *B. divergens* isolates (R87, Spanish, and Turkiye isolates) revealed sequence differences in the IG region. While the *EF-1α* gene sequences were found to be 100% identical among all isolates, the IG region length varied across the isolates, with the R87 strain containing 5 TA repeats and measuring 1380 bp, the Spanish isolate harboring 24 TA repeats with an IG region of 1418 bp, and the Turkiye isolate exhibiting 8 TA repeats with an IG region of 1386 bp (Figure 3). Overall, these findings indicate notable differences in the IG region between *B. divergens* isolates originating from different geographical regions and host sources.

Multiple sequence alignment of the IG region across the three *B. divergens* isolates further confirmed variations in TA repeat numbers and additional nucleotide substitutions. The TA repeat sequences were located around position ∼513–516 bp across all isolates but varied in number. Besides TA repeat variations, additional sequence polymorphisms were identified across the IG region (Figure 4). The results were consistent and validated using Nanopore and Sanger sequencing technologies.

**Figure 4:**
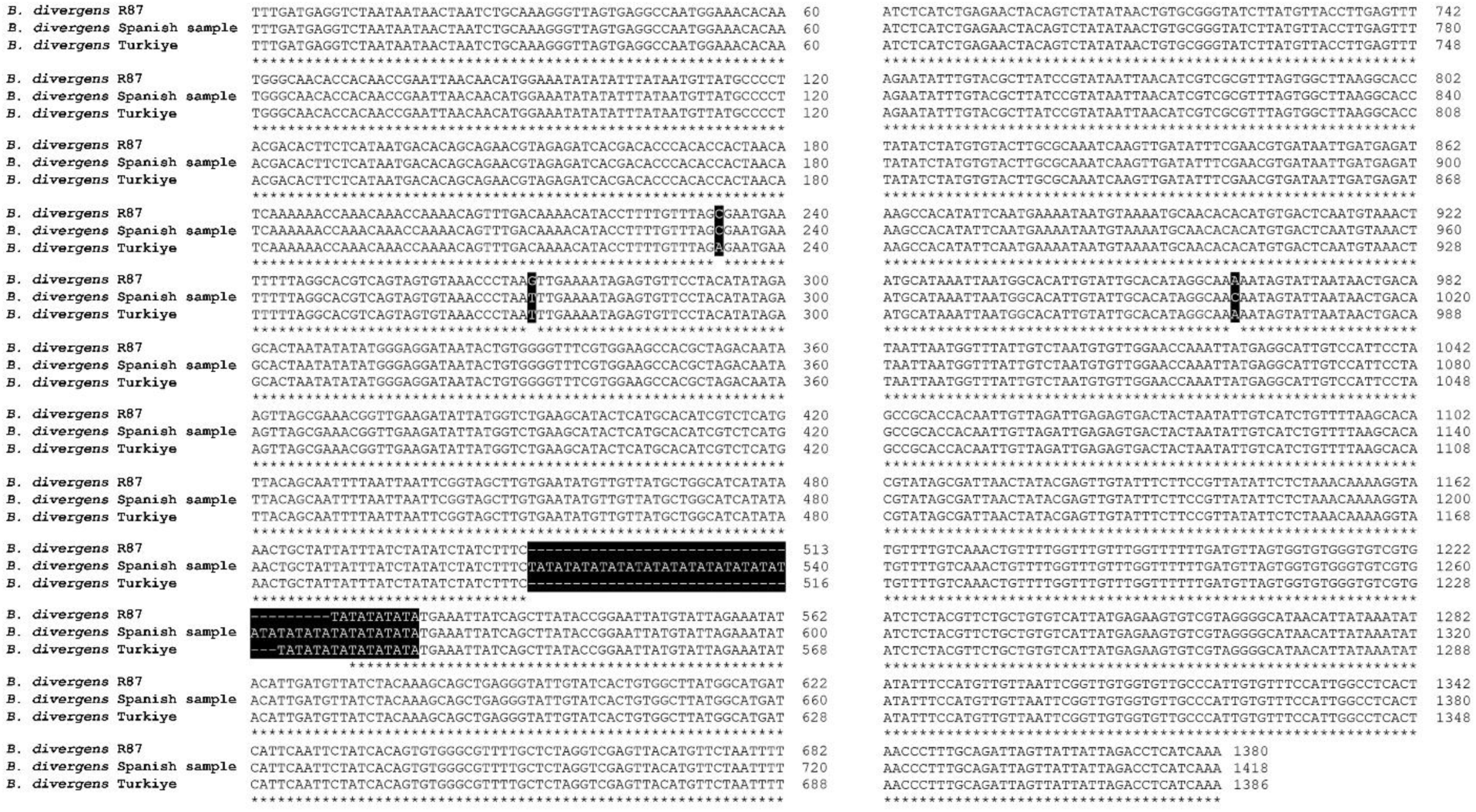
Multiple sequence alignment of the IG region within the EF-1α locus between *B. divergens* isolates. Nucleotide variations are highlighted with black boxes, while differences in TA repeat numbers are also marked. In addition to *B. divergens*, the intergenic region was examined in other zoonotic *Babesia* species, including *Babesia* MO1 and *B. duncani*. While the EF-1α locus in *B. duncani* lacked TA repeats, the *Babesia* MO1 genome contained a short stretch of 3 TA repeats. These findings indicate that the presence and expansion of TA repeats may be specific to *B. divergens* (Supplementary Figure 3).

**Figure 5:**
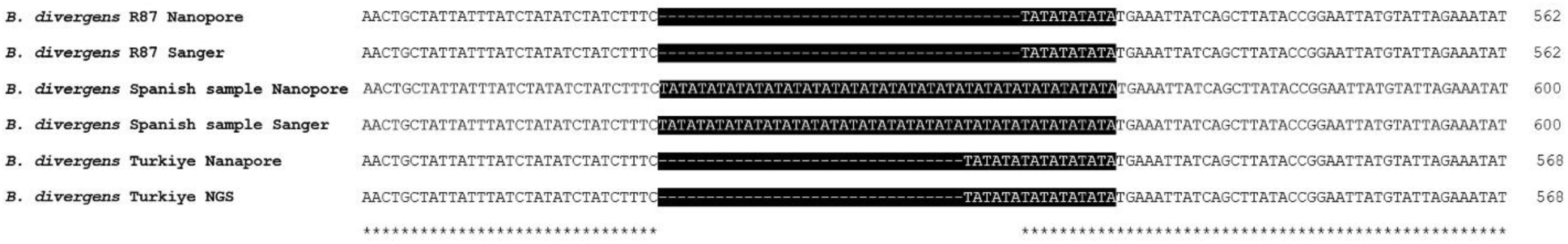
Multiple sequence alignment of the TA repeat region across *B. divergens* isolates using Nanopore, Sanger, and NGS sequencing methods. The black-highlighted regions indicate TA repeats.

### Validation of TA repeat variations in the IG region

To confirm the sequence differences in the TA repeat region among *B. divergens* isolates, a combination of PCR amplification, Sanger sequencing, and NGS approaches was employed. For the R87 isolate, the IG region was amplified using the Bdiv-Ef-For1/IGRev1 primer pair, yielding an amplicon of approximately 2.8 kb, which was subsequently validated by Sanger sequencing. The repeat structure identified in the sequencing data confirmed the presence of 5 TA repeats, consistent with initial findings. For the Turkiye isolate, the IG region was amplified in two overlapping fragments using the Bdiv-Ef-For1/Sez-For1 and Sez-For1-Rev/GlutamylR1 primer sets. These amplicons were subjected to NGS analysis, which verified the 8 TA repeats and confirmed the overall sequence integrity of the IG region. For the Spanish isolate, initial attempts to amplify the full IG region using Bdiv-Ef-For1/IGRev1 primers were unsuccessful, likely due to sequence divergence at the primer binding sites. An alternative primer pair, IGLMFR1/IGRev1, was used to amplify ∼1 kb fragment, which was validated through Sanger sequencing. The results confirmed the presence of 24 TA repeats, aligning with the Nanopore sequencing data.

## Discussion

Since the discovery of *Babesia*, the availability of transfection and gene editing methods arguably emerged as the most significant developments, after *Babesia in vitro* cultures. Transfection technologies (together with related gene editing methods), which involves the introduction of foreign genetic material into cells, has revolutionized the study of various diseases, including those caused by *Babesia* parasites [15]. This technique has enabled researchers to manipulate cellular processes, elucidate pathogenic mechanisms, and explore potential therapeutic interventions. Thus, transfection represents a crucial tool in advancing our understanding of *Babesia* infections and developing strategies for their prevention and treatment [13,17]. EF-1α-IG region contains bi-directional promoters capable of efficiently driving expression of *EF-1α* genes. Their utilization has significantly advanced transfection systems in *Babesia* [11,15]. In this study, we explored the structural and sequence variability of the EF-1α-IG regions across three *B. divergens* isolates from humans and cattle to better understand its potential role in gene regulation and parasite adaptation. The findings indicated that while the EF-1α coding sequences remain highly conserved, there is variability in the IG region, specifically in TA repeat numbers and the length of the terminal IR sequences, suggesting that these elements could contribute to transcriptional regulation and genomic stability.

The high level of conservation of the *EF-1α* gene observed in different strains of *B. divergens* and its similarity to the EF-1α protein of the two other pathogenic bovine babesiosis species, *B. bovis* and *B. bigemina*, highlights the potential use of EF-1α in subunit vaccines. These findings revealed that EF-1α protein has potential to be a suitable antigen for vaccine development, as it was previously suggested for other important protozoa such as *T. gondii* [9] and *C. parvum* [8], which are also members of the apicomplexan phylum. Additionally, the broad applicability of the vaccine in suppressing different parasitic infections was further verified by the existing cross-protective immunity against *Eimeria tenella* and *Eimeria maxima* in broiler chickens vaccinated with recombinant EF-1α protein [10]. Thus, EF-1α protein appears as a good candidate for the development of vaccines against babesiosis, taking advantage of its potential to induce cross-immunity against different species.

Comparisons of the IG regions among the three *B. divergens* isolates (R87, Spanish, and Turkiye isolates) provided novel insights into the structural organization of this region. This study challenged a previously established assumption that the IG regulatory regions of the EF-1α loci are fully conserved among *Babesia* spp. and parasite isolates, both in terms of sequence and structural organization. Despite having identical *EF-1α* gene ORFs organized in a typical head-to-head fashion, *B. divergens* isolates exhibited striking differences in their IG regions, despite sharing identical *18S rRNA* sequences. One of the most unexpected findings of this study was the variability in TA repeat numbers within the IG region. The Spanish isolate exhibited the highest number of TA repeats (n=24), while the Turkiye isolate contained 8, and the R87 strain had only 5. Given that repeat sequences in intergenic regions can affect promoter activity, transcription factor binding, and chromatin accessibility [31], these differences may influence *EF-1α* gene expression. The conserved 237 bp terminal IR region and the 153 bp intron within the untranslated region suggest that the IG region may have multiple regulatory roles. Inverted repeats (IRs) are often associated with promoter function, transcriptional regulation, while intronic sequences within untranslated regions can influence alternative splicing, mRNA stability, or post-transcriptional control. The conservation of this structure across isolates suggests its functional importance in *EF-1α* gene regulation.

As mentioned, the differences within the IG regions among *B. divergens* isolates analyzed hereby predominantly occur in the initial segment of the putative promoter regions. Although investigations into EF-1α-IG region variations across different strains remain limited, examination of these regions from *B. bovis* strains in Argentina, Mexico, and Australia suggests a notable degree of conservation among them [11]. The distinctions between *B. bovis* and *B. divergens*, despite both being etiological agents of babesiosis in cattle, are evident in their geographical distribution, different tick vector preferences, and the clinical manifestations of babesiosis [23,32]. Moreover, the broader host spectrum of *B. divergens*, including its zoonotic potential, highlights its adaptability and significance beyond the confines of bovine hosts. It is thus logical to speculate that the discrepancies observed in the EF-1α-IG region could be attributed to the differing host preferences of *B. divergens*. This is particularly noteworthy in our study where R87, Spanish, and Turkiye isolates, whose elongation loci we compared, were obtained from humans (R87, and Spanish isolates) and cattle (Turkiye) respectively. Surprisingly, the EF-1α-IG region from parasites that infect humans presented differences that could also be related to virulence or adaptation to laboratory conditions. Such findings raise intriguing questions regarding factors involved in parasite evolution, and the potential influence of host-specific factors on the genetic makeup and regulatory mechanisms of *Babesia* species, warranting further investigation into the intricate interplay between pathogen genetics and host environments. As they conflict with a previously established paradigm, the substantial differences found in the IG regions of the *EF-1α* genes between three different strains of *B. divergens*, as confirmed by *18S rRNA gene* analysis are found for the first time among piroplasmid parasites. Moreover, as previously noted, these differences may reflect important adaptive events affecting the development of the parasites among distinct invertebrate and/or vertebrate hosts.

Surprisingly, upon expanding our comparative analysis of the EF-1α IG regions found in other zoonotic *Babesia* species, *B. duncani* and *Babesia* MO1, striking differences were also noted in their EF-1α IG regions. *Babesia duncani*, on the other hand, contained no TA repeats, while only 3 TA motifs were present in the MO1 isolate. This suggests that TA repeat expansion within the EF-1α-IG region may represent a species-specific regulatory feature rather than a common trait of all zoonotic *Babesia*. Further comparative studies across more isolates will be essential to determine whether these differences contribute to host tropism, gene expression variability, or virulence mechanisms.

## Conclusion

This study provides new insights into the structural variability of the EF-1α-IG region in *B. divergens* isolates from different hosts, revealing differences in TA repeat numbers that may contribute to gene regulation and parasite adaptation. While our findings highlight the potential regulatory role of this region, functional validation through promoter activity assays, gene expression analysis, and transfection studies is still needed to determine its precise impact on EF-1α expression. Given the zoonotic nature of *B. divergens*, future research should explore whether similar variations in the IG region exist in *B. divergens* isolates from cattle and humans. Additionally, the gerbil model could be used to investigate potential intergenic region variations across different vertebrate hosts to better understand the role of this region in host adaptation. Further investigation of the functional implication of these variations may provide valuable insights into transcriptional control across diverse parasite strains, provide valuable information on the mechanisms driving *B. divergens* evolution and improve transfection strategies for functional genomic studies. Overall, this study lays the groundwork for future research into gene regulation and host-pathogen interactions in zoonotic *Babesia* species.

## Methods

### *Babesia divergens* -Rouen 87 Strain (R87)

*Babesia divergens* R87 is a reference strain originally isolated from a human case of babesiosis in France [33]. This strain has been widely used in molecular and genomic studies due to its well-characterized genome sequence [27]. The EF-1α locus data from this strain were utilized in this study, obtained from the *B. divergens* genome assembly described by [27].

### Babesia divergens-Spanish sample

*Babesia divergens* Spanish strain was isolate from a blood sample of elderly patient from Asturias, Spain who suffers a fulminant fatal babesiosis [28]. The blood sample was cultured in vitro in human A+ RBCs and RPMI 1640 (Gibco-Life Technology, 11875093) supplemented with 10% human serum (Gibco, 11021037), 7.5% (w/v) sodium bicarbonate solution (Lonza Group Ltd, Basel, Switzerland, 144-55-8), and 100 μmol/L hypoxanthine (Sigma-Aldrich Corporation, St Louis, MO, H9377) at a pH of 7.3. Cells were cultured at 37°C in a humidified atmosphere of 5% CO2. The culture medium was replaced every 24 h, and parasitemia was monitored by examining Giemsa-stained blood smears by a light microscope.

### Babesia divergens-Turkiye

*Babesia divergens*-Turkiye isolate was obtained from a 3-year-old cow exhibiting severe babesiosis symptoms, including high fever, anemia, and hemoglobinuria, in Bartin province of Turkiye in 2023. Microscopic examination revealed a parasitemia level of 1.5%. Following the collection of blood stabilate from the animal, treatment was administered using imidocarb dipropionate [34]. To determine the species of *Babesia*, Reverse Line Blot (RLB) analysis was conducted. Total DNA extraction from blood samples was performed using the PureLink™ Genomic DNA Mini Kit (Invitrogen Corporation, Carlsbad, United States) according to the manufacturer’s instructions. The hypervariable V4 region of the piroplasm *18S rRNA* gene was then amplified using primers RLB-F2 and RLB-R2-biotin [35] specifically for the RLB assay. RLB was carried out on the PCR product, following the previously described method [36]. After amplification, 20 µL of all PCR products obtained from each DNA sample were diluted to a final volume of 150 µL with 2X SSPE/0.1% SDS buffer. For RLB hybridization, the samples were heated at 95–100°C for 10 minutes in a Thermal Cycler and rapidly cooled on ice. Subsequently, the PCR products were hybridized with probes specific to the genera and species of *Babesia* and *Theileria*, which were linked to an RLB membrane [36].

### Babesia MO1 and Babesia duncani

In this study, we used the EF-1α locus data of these species obtained from the *Babesia* MO1 [37] and *B. duncani* [38,39] assembly. Specifically, Bioproject PRJNA1032622 for *Babesia MO1* and Bioproject PRJNA821606 and locus OL804102 for *B. duncani.* To assess the presence of TA repeats in *Babesia* MO1 and *B. duncani*, we retrieved the EF-1α locus from *Babesia* MO1 and *B. duncani* genome assemblies (PRJNA1032622 and PRJNA821606, respectively). The intergenic regions were identified by locating the flanking genes (*ribonucleoside diphosphate reductase* and *glutamyl tRNA synthetase*) and were subsequently aligned to detect potential tandem repeat regions. The presence or absence of TA repeats was manually inspected by multiple sequence alignment.

### Nanopore sequencing

The total DNA from *Babesia* samples was used to prepare Oxford Nanopore libraries with the SQK-LSK114 Ligation Sequencing Kit for 24 h in the MinION Platform with an R10.4 flow cell following the manufacturer’s protocol (Oxford Nanopore Technologies, Oxford, United Kingdom). The base calling was performed using the software Dorado v0.8.3 with the duplex option. The sequencing and basecalling was carried out at the Unidad Universitaria de Secuenciación Masiva y Bioinformática, Instituto de Biotecnología, Universidad Nacional Autónoma de México (Cuernavaca, Mexico).

### Designing of primers for EF-1α locus analysis and PCR

Primers were designed utilizing sequences from *Ribonucleoside diphosphate reductase* (Bdiv_023390c) and *Glutamyl tRNA* (Bdiv_023400) genes obtained from the Piroplasma DB (https://piroplasmadb.org) and nanopore sequencing data to amplify the EF-1α locus. All primers were designed using Primer Quest™ Tool (https://www.idtdna.com/pages/tools/primerquest) (Supplementary Table 1, and Supplementary Figure 4). PCR amplification was carried out using Phusion® High-Fidelity PCR Master Mix with GC Buffer (M0532S; NEB). The PCR was performed in a total reaction volume of 20 μL containing 10 μL of 2X Phusion Master Mix, 1 μL of each forward and reverse primer, 1 μL of template DNA, 7 μL of nuclease-free water.

### Illumina sequencing and *de novo* assembly

Sequencing library was prepared using Nextera XT DNA Library Preparation Kit and sequencing was performed by Illumina Miseq platform as paired end (PE) 2x150 bases reads. Raw NGS reads (FASTQ) were quality checked by FASTQC [40] and trimmed by Trimmomatic v0.32 [41]. Demultiplexing and low-quality read filtering were performed via CLC Genomics Workbench (Qiagen, Hilden, Germany). The *de novo* assembly constructed by CLC Genomic’s de novo assembly module with the parameters of minimum contig size as 200 bp, mismatch cost is 2, insert cost is 3, deletion cost is 3, length fraction is 0.5, similarity fraction is 0.8, and paired read input as minimum distance of 180.

### Sanger sequencing

To validate the assembled EF-1α-IG region, Sanger sequencing was performed on PCR amplified IG regions from the *B. divergens* isolates. PCR products were purified using the mi-Gel Extraction Kit (Metabion international AG, Steinkirchen Germany) and sequenced using an ABI PRISM 3730XL DNA Analyzer (Applied Biosystems, San Francisco, CA, USA). The resulting chromatograms were analyzed using manually inspected, corrected, and edited using Chromas Pro program (McCarthy, Queensland, Australia) and Laser Gene 12.1 program (DNAStar, Madison, WI, USA), and sequences were aligned against the de novo assembled IG region to confirm accuracy and resolve potential misassembles.

### Phylogenetic analysis

Phylogenetic relationships were inferred using Molecular Evolutionary Genetics Analysis (MEGA 11) [42]. The maximum likelihood (ML) method was employed to construct phylogenetic trees based on EF-1α protein sequences and *18S rRNA* gene sequences from various piroplasm species. Bootstrap support values were calculated with 1,000 replicates to assess tree robustness.

## Data availability

All data is provided within the manuscript or supplementary information files.

## Ethical statement

*B. divergens* DNA samples from the cattle were obtained as part of routine veterinary diagnostic procedures from a clinically infected animal. No experimental infection or additional intervention was performed for research purposes. Therefore, Institutional Animal Care and Use Committee (IACUC) or Ethics Committee approval was not required. The *B. divergens* Spanish strain was obtained as part of routine diagnostic procedures, no additional ethics approval or informed consent from hospitals or patients was required. Therefore, our study does not involve human subjects, identifiable human tissue, or personal data, in accordance with the Declaration of Helsinki.

## Acknowledgements

We thank “Centro de Transfusiones de la Comunidad de Madrid”, which provided the human A+ blood from healthy volunteer donors. This work was funded by grants from Health Institute Carlos III (ISCIII; PI20CIII-00037 to E.M. and L.G.M) and Comunidad Autónoma de Madrid (CAM; TEC-2024/BIO-66/SALAINDEC-CM to EM and L.G.M.).

## Author Contrubitions

Conceptualization: SO, ASF, EM and CES; Methodology and experiments: OS, LMG, HA, AGI, RGB, MA and RG; Sequencing and bioinformatic analysis: SO, RG, LMG and ASF, Formal analysis and data interpretation SO, EM, ASF and LMG; Writing original draft SO; Writing, reviewing and editing ASF, EM, LMG. All authors read and approved of the final manuscript.

## Ethics statement

*Babesia divergens* Spanish isolate were cultured using human A+ blood obtained from healthy volunteer donors. The blood was sourced from the Blood Transfusion Center of Madrid, Spain, adhering to approved protocols and in compliance with the relevant institutional guidelines and regulations. *Babesia divergens* Turkiye isolate was obtained from a clinically diagnosed babesiosis case in a cow. The sample was submitted for diagnostic purposes to the Department of Parasitology, Faculty of Veterinary Medicine, Fırat University. The study was conducted in accordance with institutional guidelines and regulations.

## Additional information

### Supplementary Information

Supplementary Figure 1: Percentage of identity of amino acid sequence of EF-1α among different piroplasm species.

Supplementary Figure 2: Comparison of EF-1α protein sequences among *B. divergens* isolates (R87, Spanish sample, and Turkiye isolates) and various piroplasm species. Three conserved EF-1α regions with known Guanosine 5’-triphosphate (GTP)-Binding Site Function (GTP BS 1, 2, and 3) highlighted in boxes.

Supplementary Figure 3: Comparative alignment of the EF-1α-IG region across zoonotic *Babesia* species. The TA repeat motifs (highlighted in black) show notable variability among *B. divergens* isolates (R87, Spanish, and Turkiye), while *Babesia* MO1 contains only 3 TA motifs, and *B. duncani* lacks any TA repeats.

Supplementary Figure 4: Schematic representation of the *B. divergens* EF-1α locus showing the primers used for IG region amplification. IR (inverted repeat) and IG (intergenic region) are highlighted, with primer positions indicated by arrows.

## References

1. Slobin, L. I. (1980). The Role of Eucaryotic Elongation Factor Tu in Protein Synthesis: The Measurement of the Elongation Factor Tu Content of Rabbit Reticulocytes and Other Mammalian Cells by a Sensitive Radioimmunoassay. European Journal of Biochemistry 110, 555–563. doi: 10.1111/j.1432-1033.1980.tb04898.x

2. Merrick, W. C. (1992). Mechanism and regulation of eukaryotic protein synthesis. Microbiol Rev 56, 291–315. doi: 10.1128/mr.56.2.291-315.1992

3. Sasikumar, A. N., Perez, W. B., and Kinzy, T. G. (2012). The many roles of the eukaryotic elongation factor 1 complex. WIREs RNA 3, 543–555. doi: 10.1002/wrna.1118

4. Riis, B., Rattan, S. I., Clark, B. F., and Merrick, W. C. (1990). Eukaryotic protein elongation factors. Trends in biochemical sciences 15, 420–424.

5. Stapulionis, R., Kolli, S., and Deutscher, M. P. (1997). Efficient mammalian protein synthesis requires an intact F-actin system. Journal of Biological Chemistry 272, 24980–24986.

6. Chen, E., Proestou, G., Bourbeau, D., and Wang, E. (2000). Rapid up-regulation of peptide elongation factor EF-1α protein levels is an immediate early event during oxidative stress-induced apoptosis. Experimental cell research 259, 140–148.

7. Lamberti, A., Longo, O., Marra, M., Tagliaferri, P., Bismuto, E., Fiengo, A., et al. (2007). C-Raf antagonizes apoptosis induced by IFN-α in human lung cancer cells by phosphorylation and increase of the intracellular content of elongation factor 1A. Cell Death & Differentiation 14, 952– 962.

8. Matsubayashi, M., Teramoto-Kimata, I., Uni, S., Lillehoj, H. S., Matsuda, H., Furuya, M., et al. (2013). Elongation factor-1α is a novel protein associated with host cell invasion and a potential protective antigen of Cryptosporidium parvum. Journal of Biological Chemistry 288, 34111– 34120.

9. Wang, S., Zhang, Z., Wang, Y., Gadahi, J. A., Xu, L., Yan, R., et al. (2017). Toxoplasma gondii elongation factor 1-alpha (TgEF-1α) is a novel vaccine candidate antigen against toxoplasmosis. Frontiers in microbiology 8, 168.

10. Lin, R.-Q., Lillehoj, H. S., Lee, S. K., Oh, S., Panebra, A., and Lillehoj, E. P. (2017). Vaccination with Eimeria tenella elongation factor-1α recombinant protein induces protective immunity against E. tenella and E. maxima infections. Veterinary Parasitology 243, 79–84.

11. Suarez, C. E., Norimine, J., Lacy, P., and McElwain, T. F. (2006). Characterization and gene expression of Babesia bovis elongation factor-1α. International journal for parasitology 36, 965– 973.

12. Silva, M. G., Knowles, D. P., Mazuz, M. L., Cooke, B. M., and Suarez, C. E. (2018). Stable transformation of Babesia bigemina and Babesia bovis using a single transfection plasmid. Scientific reports 8, 6096.

13. Hakimi, H., Asada, M., and Kawazu, S. (2021). Recent advances in molecular genetic tools for babesia. Veterinary Sciences 8, 222.

14. Vinkenoog, R., Sperança, M. A., van Breemen, O., Ramesar, J., Williamson, D. H., Ross-MacDonald, P. B., et al. (1998). Malaria parasites contain two identical copies of an elongation factor 1 alpha gene. Molecular and biochemical parasitology 94, 1–12.

15. Suarez, C. E., and McElwain, T. F. (2010). Transfection systems for Babesia bovis: a review of methods for the transient and stable expression of exogenous genes. Veterinary parasitology 167, 205–215.

16. Fernandez-Becerra, C., Azevedo, M. de, Yamamoto, M. M., and Portillo, H. del (2003). Plasmodium falciparum: new vector with bi-directional promoter activity to stably express trans genes. Available at: https://www.cabidigitallibrary.org/doi/full/10.5555/20033124083 (Accessed March 13, 2025).

17. Suarez, C. E., Alzan, H. F., Silva, M. G., Rathinasamy, V., Poole, W. A., and Cooke, B. M. (2019). Unravelling the cellular and molecular pathogenesis of bovine babesiosis: is the sky the limit? International journal for parasitology 49, 183–197.

18. Johnson, W. C., Hussein, H. E., Capelli-Peixoto, J., Laughery, J. M., Taus, N. S., Suarez, C. E., et al. (2022). A transfected Babesia bovis parasite line expressing eGFP is able to complete the full life cycle of the parasite in mammalian and tick hosts. Pathogens 11, 623.

19. Laughery, J. M., Knowles, D. P., Schneider, D. A., Bastos, R. G., McElwain, T. F., and Suarez, C. E. (2014). Targeted surface expression of an exogenous antigen in stably transfected Babesia bovis. PLoS One 9, e97890.

20. Oldiges, D. P., Laughery, J. M., Tagliari, N. J., Leite Filho, R. V., Davis, W. C., da Silva Vaz Jr, I., et al. (2016). Transfected Babesia bovis expressing a tick GST as a live vector vaccine. PLoS neglected tropical diseases 10, e0005152.

21. Suarez, C. E., Bishop, R. P., Alzan, H. F., Poole, W. A., and Cooke, B. M. (2017). Advances in the application of genetic manipulation methods to apicomplexan parasites. International journal for parasitology 47, 701–710.

22. Mazuz, M. L., Laughery, J. M., Lebovitz, B., Yasur-Landau, D., Rot, A., Bastos, R. G., et al. (2021). Experimental infection of calves with transfected attenuated Babesia bovis expressing the Rhipicephalus microplus Bm86 Antigen and eGFP Marker: Preliminary studies towards a dual anti-tick/Babesia vaccine. Pathogens 10, 135.

23. Ozubek, S., Bastos, R. G., Alzan, H. F., Inci, A., Aktas, M., and Suarez, C. E. (2020). Bovine babesiosis in Turkey: Impact, current gaps, and opportunities for intervention. Pathogens 9, 1041.

24. Hildebrandt, A., Zintl, A., Montero, E., Hunfeld, K.-P., and Gray, J. (2021). Human babesiosis in Europe. Pathogens 10, 1165.

25. Cubillos, E. F., Snebergerova, P., Borsodi, S., Reichensdorferova, D., Levytska, V., Asada, M., et al. (2023). Establishment of a stable transfection and gene targeting system in Babesia divergens. Frontiers in Cellular and Infection Microbiology 13. Available at: https://www.ncbi.nlm.nih.gov/pmc/articles/PMC10753763/ (Accessed April 30, 2024).

26. Cuesta, I., González, L. M., Estrada, K., Grande, R., Zaballos, Á., Lobo, C. A., et al. (2014). High-Quality Draft Genome Sequence of Babesia divergens, the Etiological Agent of Cattle and Human Babesiosis. Genome Announc 2, e01194–14. doi: 10.1128/genomeA.01194-14

27. Gonzalez, L. M., Estrada, K., Grande, R., Jiménez-Jacinto, V., Vega-Alvarado, L., Sevilla, E., et al. (2019). Comparative and functional genomics of the protozoan parasite Babesia divergens highlighting the invasion and egress processes. PLoS neglected tropical diseases 13, e0007680.

28. Asensi, V., González, L. M., Fernández-Suárez, J., Sevilla, E., Navascués, R. Á., Suárez, M. L., et al. (2018). A fatal case of Babesia divergens infection in Northwestern Spain. Ticks and tick-borne diseases 9, 730–734.

29. Le, S. Q., and Gascuel, O. (2008). An improved general amino acid replacement matrix. Molecular biology and evolution 25, 1307–1320.

30. Tamura, K., and Nei, M. (1993). Estimation of the number of nucleotide substitutions in the control region of mitochondrial DNA in humans and chimpanzees. Molecular biology and evolution 10, 512–526.

31. Quilez, J., Guilmatre, A., Garg, P., Highnam, G., Gymrek, M., Erlich, Y., et al. (2016). Polymorphic tandem repeats within gene promoters act as modifiers of gene expression and DNA methylation in humans. Nucleic acids research 44, 3750–3762.

32. Schnittger, L., Ganzinelli, S., Bhoora, R., Omondi, D., Nijhof, A. M., and Florin-Christensen, M. (2022). The Piroplasmida Babesia, Cytauxzoon, and Theileria in farm and companion animals: Species compilation, molecular phylogeny, and evolutionary insights. Parasitology research 121, 1207–1245.

33. Gorenflot, A., Brasseur, P., Precigout, E., L’Hostis, M., Marchand, A., and Schrevel, J. (1991). Cytological and immunological responses toBabesia divergens in different hosts: Ox, gerbil, man. Parasitol Res 77, 3–12. doi: 10.1007/BF00934377

35. Georges, K., Loria, G. R., Riili, S., Greco, A., Caracappa, S., Jongejan, F., et al. (2001). Detection of haemoparasites in cattle by reverse line blot hybridisation with a note on the distribution of ticks in Sicily. Veterinary parasitology 99, 273–286.

34. Mosqueda, J., Olvera-Ramírez, A., Aguilar-Tipacamú, G., and Cantó, G. (2012). Current Advances in Detection and Treatment of Babesiosis. Curr Med Chem 19, 1504–1518. doi: 10.2174/092986712799828355

36. Aktas, M., and Ozubek, S. (2015). Molecular and parasitological survey of bovine piroplasms in the Black Sea region, including the first report of babesiosis associated with Babesia divergens in Turkey. Journal of medical entomology 52, 1344–1350.

37. Singh, P., Vydyam, P., Fang, T., Estrada, K., Gonzalez, L. M., Grande, R., et al. (2024). Insights into the evolution, virulence and speciation of Babesia MO1 and Babesia divergens through multiomics analyses. Emerging Microbes & Infections 13, 2386136. doi: 10.1080/22221751.2024.2386136

38. Wang, S., Li, D., Chen, F., Jiang, W., Luo, W., Zhu, G., et al. (2022). Establishment of a transient and stable transfection system for babesia duncani using a homologous recombination strategy. Frontiers in Cellular and Infection Microbiology 12, 844498.

39. Singh, P., Lonardi, S., Liang, Q., Vydyam, P., Khabirova, E., Fang, T., et al. (2023). Babesia duncani multi-omics identifies virulence factors and drug targets. Nature microbiology 8, 845– 859.

40. Bioinformatics, B. (2018). Babraham Bioinformatics-FastQC a quality control tool for high throughput sequence data.

41. Bolger, A. M., Lohse, M., and Usadel, B. (2014). Trimmomatic: a flexible trimmer for Illumina sequence data. Bioinformatics 30, 2114–2120.

42. Tamura, K., Stecher, G., and Kumar, S. (2021). MEGA11: molecular evolutionary genetics analysis version 11. Molecular biology and evolution 38, 3022–3027.

