## Supplementary material for "Comparative analysis of the EF-1α Intergenic Region in *Babesia divergens* isolates: Insights into TA Repeat Variation and Potential Regulatory Implications": suppl material: Supplementary Table 1.docx

**Supplementary Table 1.** Detailed information on the primers used in PCR assays.

| **Primer** | **Sequence (5’-3’)** |
| --- | --- |
| **Bdiv-Ef-For1** | TTCCCAGTCCTTCATATC |
| **Sez-For1** | CCGTTTCGCTAACTTATT |
| **Sez-For1-Rev** | AATAAGTTAGCGAAACGG |
| **IGFor1** | ATCTGAGAACTACAGTCTAT |
| **IGRev1** | TCTCATCAATTATCACGTTC |
| **IGLMFR1** | CCACTGTCGACGTGGCCGATAACG |
| **GlutamylR1** | GTGATCCACTGACTTCT |
