## Supplementary figures and images for "Comparative analysis of the EF-1α Intergenic Region in *Babesia divergens* isolates: Insights into TA Repeat Variation and Potential Regulatory Implications"

### Supplementary Figure 1.jpg

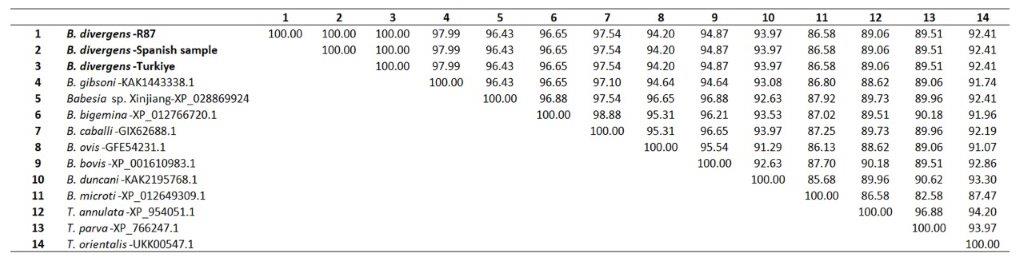

### Supplementary Figure 2.jpg

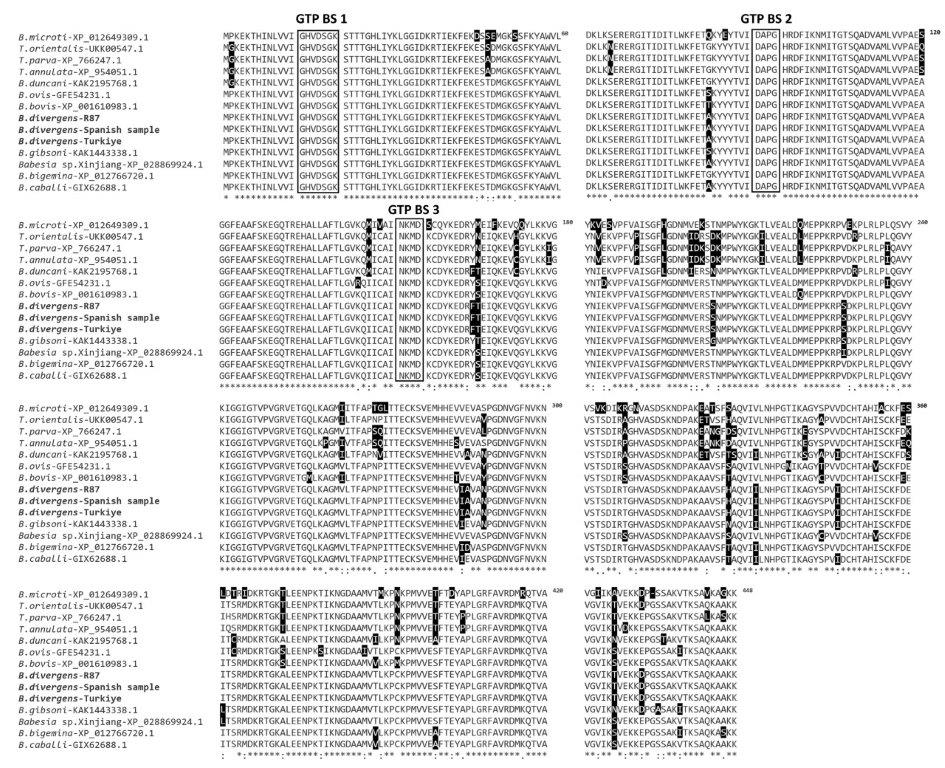

### Supplementary Figure 3.jpg

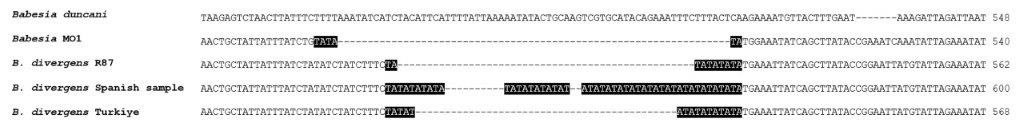

### Supplementary Figure 4.jpg

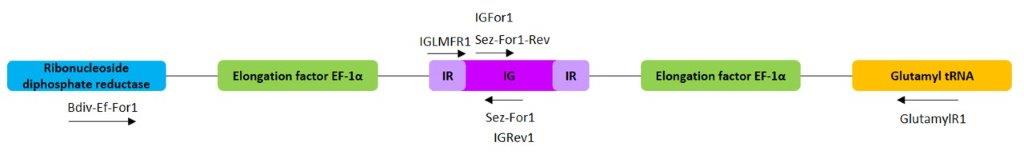
